# Integrative analysis of proteome and transcriptome dynamics during *Bacillus subtilis* spore revival

**DOI:** 10.1101/682872

**Authors:** Bhagyashree Swarge, Martijs Jonker, Wishwas Abhyankar, Huub Hoefsloot, Chris G. de Koster, Stanley Brul, Leo J. de Koning

## Abstract

*Bacillus subtilis* forms highly resistant, metabolically inactive dormant spores upon nutrient limitation. These endospores pose challenges to the food and medical sectors. Spores reactivate their metabolism upon contact with germinants and develop into vegetative cells. The activation of the molecular machinery that triggers the progress of germination and spore outgrowth is still unsettled. To gain further insight in spore germination and outgrowth processes, the transcriptome and proteome changeover during spore germination and outgrowth to vegetative cells, was analysed. *B. subtilis* transcriptome analysis allow us to trace the different functional groups of genes expressed. For each time-point sample, the change in the spore proteome was quantitatively monitored relative to the reference proteome of ^15^N metabolically labelled vegetative cells. We observed until the phase transition, i.e. completion of germination, no significant change in the proteome. We have identified 36 transcripts present abundantly in the dormant spores. This number is in close agreement with the previous findings. These transcripts mainly belong to the genes encoding small acid soluble proteins (*sspE, sspO, sspI, sspK, sspF*) and proteins with uncharacterized functions. We observed in total 3152 differentially expressed genes, but ‘only’ 323 differentially expressed proteins (total 451 proteins identified and quantified). Our data shows that 173 proteins from dormant spores, both spore unique proteins and protein shared with vegetative cells, are lost during the phase transitioning period. This loss is in addition to the active protein degradation, undertaken by the spore proteases such as Gpr, as germination and outgrowth proceeds. Further analysis is required to functionally interpret the observed protein loss. The observed diverse timing of the synthesis of different protein sets reveals a putative core-strategy of the revival of ‘life’ starting from the *B. subtilis* spore.

## 1. Introduction

Nutrient fluctuations in the environment, lead to a switch between the vegetative state and the dormant state in *Bacillus subtilis*. This ability to transform into a metabolically dormant, heat resistant endospore and revert from this latent state to proliferative vegetative cell is studied in bacilli and clostridia since few decades. The molecular mechanisms controlling these two processes are well studied, and its constituting proteins have been to a large extent identified through genetic analysis. Their actual biochemical structure and function are, however, in many ways only beginning to be appreciated^1-4^. Through the germinant sensing and calcium dipicolinic acid transport signal transduction system, *B. subtilis* channels information related to external stimuli to an internal response, enabling the above mentioned morphogenetic transformation. The dormant spores harbour the transcriptional and translational machinery that might facilitates the early stages of outgrowth^5^. The protein cargo that transfers from progenitor cell to the spore, the ‘tool-kit for life’, is key to the spore’s memory^6^ which in turn modulates its germination response^7^. Thus, clearly an in-depth research on pre-existing transcripts and proteins in spore revival is important to understand long-term stress survival. Recently, a few glycolytic enzymes and alanine dehydrogenase present in dormant spores have been identified to make up the spore’s phenotypic memory in *Bacillus anthracis*^7^.

After committing to germination, spores release monovalent cations and CaDPA followed by cortex hydrolysis by the cortex lytic enzymes. This leads to complete hydration of the spore core thus completing germination and leading to the reactivation of the molecular machinery for the emerging vegetative cell. Stores of 3PGA in the spore’s core are utilized to generate ATP^2,8,9^ initiating macromolecular synthesis. A study monitoring the temporal expression of mRNA during spore germination and outgrowth has demonstrated the systematic onset of transcription^1,4^ whereas a detailed analysis of reviving spores has provided information on newly synthesized proteins of *Bacillus subtilis* spores including a putative set of enzymes that are crucial for the onset of protein synthesis^3,10^. One particular aspect of spore revival, whether protein synthesis is required for germination to occur or not, is topic of much recent debate ^9,11,12^.

Here we report, in contrast to previous studies where ‘only’ newly synthesized proteins were revealed, on the proteome ‘turnover’ during spore germination and outgrowth to vegetative cells. The data is linked to the changing transcriptome to uncover the dynamic relationship between mRNA and protein levels in reviving *B. subtilis* spores. With ^15^N metabolic labelling and a SILAC approach for relative protein quantification, a unique detailed time-series displaying the protein profiles at different stages of spore revival is linked to a microarray-analysis derived transcriptome. Our data show that, protein synthesis does not occur in a phase bright spore. The phase transition seems to coincide with both transcription and translation following the initial degradation of spore protein repositories. Specific functional protein modules were inferred that might play crucial regulatory roles in progressing through the various phases of *Bacillus subtilis* spore germination and outgrowth.

## 2. Material and methods

### 2.1 Growth conditions, sporulation

The *B. subtilis* strains PY79 and PB2 were used in this study. Growth conditions for sporulation are described elsewhere^5^. Mature spores (96 hrs) were purified as described previously^13^. For ^15^N metabolic labelling, *B. subtilis* PY79 cells were grown as described previously^5^. Briefly, NH_4_Cl supplied in MOPS medium was replaced with ^15^NH_4_Cl. Vegetative cells were harvested during the exponential growth phase and washed two times with phosphate buffer saline pH 7.5.

### 2.2 Germination assay

Purified spores were heat activated (HA) (70°C for 30 minutes) prior to germination. The spores were suspended in MOPS liquid minimal medium (pH 7.4) supplemented with a mixture of AGFK (10mM L-asparagine, 10mM D-glucose, 1mM D-fructose, 1mM KCl) and 10mM L-alanine. Throughout the spore revival, samples were drawn at regular time intervals for further analysis (**Figure 1**). For statistical purposes, three replicates were taken for each time point for both transcriptional and proteomic analyses. Each replicate originated from a different batch of spores.

**Figure 1.**
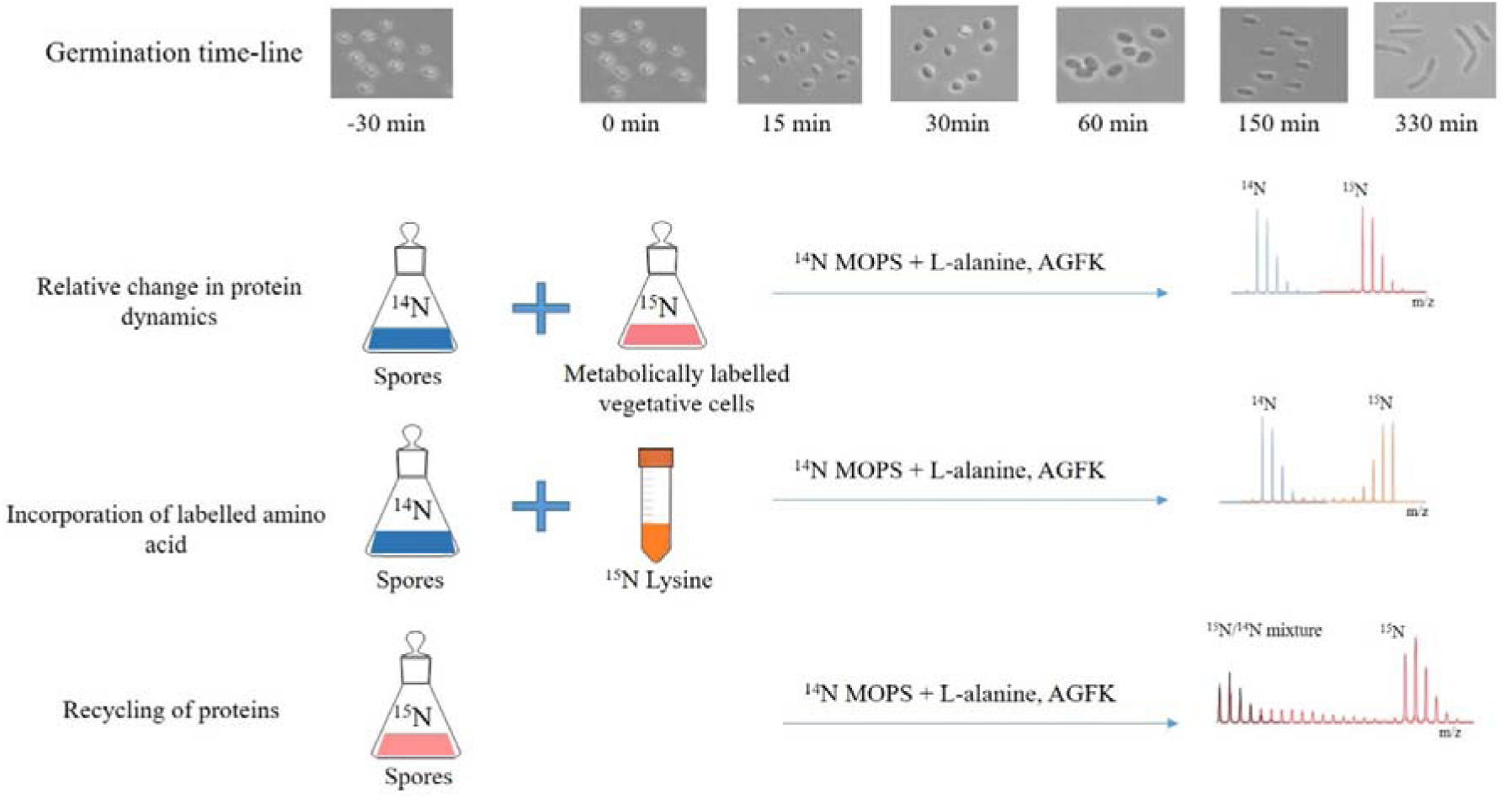
Germination timeline. **A)** Morphological changes during spore germination and outgrowth were investigated by microscopic analysis. Samples were harvested at different time points during spore revival. **B)** Changes in spore proteins were analysed relative to ^15^N-metabolically labelled vegetative cells. This approach was used to study spore germination timeline unless otherwise stated. **C)** Incorporation of ^15^N_2_^13^C_6_Lysine in protein, an indication of protein synthesis was monitored during early and later stages of outgrowth. **D)** Amino acid turnover throughout spore revival was monitored by germinating ^15^N-labelled spore in ^14^N-minimal medium.

### 2.3 Microscopy

Light microscopy was used with a Zeiss Axiostar plus microscope (CP-achromat 100x/1,25 oil objective), in combination with a Lumenera INFINITY2-2 digital charge coupled device (CCD).

### 2.4 Transcriptome analysis

For transcriptome analysis of *B. subtilis* PY79, dormant spores before HA (t = −30 min), during HA (t = −15 min) after HA (t = 0 min) were also analysed in addition to the germinating spores (t = 15, 30, 60, 150, 330 min).

#### 2.4.1 RNA isolation

A variation of the protocol detailed in the RNeasy MinElute Cleanup kit from Qiagen was used for all extractions. Briefly, the spores sampled at given time points were centrifuged at 10 000 rpm for 5 minutes at 0°C and the supernatant was discarded. RLT buffer (with 10 μl ß-Mercaptoethanol/ml of RLT buffer) was added to each pellet and the pellet was transferred to a 2 ml screw cap tube. Spore lysis was achieved with a Precellys24 homogenizer and Zirconium beads through 7 cycles at 6000 rpm for 20 seconds and the samples were kept on ice for 2 min in between each cycle. The lysate was centrifuged at 15 000 rpm for 2 min and the supernatant was transferred to a new tube. This last step was repeated in order to ensure the removal of all cell debris/spore material. An equal volume of ice-cold 70% ethanol was added and the contents were mixed by pipetting. Further purified RNA was recovered using RNeasy spin columns according to the manufacturer’s instructions. In the final step, RNA was eluted in 50 μl RNase-free water. Subsequent to RNA isolation, DNA contamination was removed using the Turbo DNA-free kit (Ambion Inc., Applied Biosystems, USA) as per manufacturer’s guidelines. Purity and quantity of isolated RNA were checked spectrophotometrically on a NanoDrop spectrophotometer (ND-1000, Isogen Life Science, NL), while structural integrity of RNA was checked with gel electrophoresis on 1% agarose gels.

#### 2.4.2 Microarray analysis

Per sample, 100 ng of total RNA was combined with Array Control RNA Spikes (Ambion) and labelled using the Low Input Quick Amp WT Labelling kit (Agilent) according to the manufacturer’s instructions. Each hybridization mixture was made up from 1.1 μg Test (Cy3) and 1.1 μg Reference (Cy5) sample (pool of all samples). The samples were dried and 1.98 μl water was added. The hybridization cocktail was made according to the manufacturer’s instructions (NimbleGen Arrays User’s Guide – Gene Expression Arrays Version 5.0, Roche NimbleGen). Then, 7.2 μl from this mix was added to each sample. The samples were incubated for 5 min at 65°C and 5 min at 42°C prior to loading. Hybridization samples were loaded onto a 12×135K microarray custom designed against *Bacillus subtilis* (Roche NimbleGen). Microarrays were hybridized for 20 hours at 42°C with the NimbleGen Hybridization System (Roche NimbleGen). Afterwards, the slides were washed according to the NimbleGen Arrays User’s Guide – Gene Expression Arrays Version 6.0 and scanned with an Agilent DNA microarray scanner G2565CA (Agilent Technologies). Feature extraction was performed with NimbleScan v2.6 (Roche NimbleGen) by the MAD (University of Amsterdam, The Netherlands).

#### 2.4.3 Normalisation and statistical analysis

The microarray data were analysed using the R statistical language (https://cran.r-project.org/) with packages made available by the Bioconductor project (https://www.bioconductor.org/). All slides were subjected to a set of quality control checks, such as visual inspection of the scans, examining the consistency among the replicated samples by principal components analysis, testing for consistent performance of the labelling dyes, and visual inspection of pre- and post-normalized data with box plots and RI plots. After log_2_ transformation, the data was normalized using a loess smoothing procedure, based on the spikes. Log_2_ ratios were calculated for each feature and gene expression values were calculated using the median polish algorithm^14^. The normalized data was statistically analysed for differential gene expression using a mixed linear model with coefficients for Batch (random), and Time (fixed)^15,16^. A contrast analysis was applied to compare the consecutive time points. The Fs test statistic^17^ was used for hypothesis testing and the resulting *p-*values were corrected for false discoveries according to^18^.

### 2.5. Proteome Analyses

#### 2.5.1. Approach I: ^15^N-metabolic labelling for relative quantification

Samples for dormant spores at t = - 30 minutes, for spores after heat activation and adding germinants at t= 0, and during germination at t =15, 30, 45, 60, 90, 150, 210 and 330 minutes were harvested. The harvested *B. subtilis* PY79 ^14^N-spores at each time point were mixed in a 1:1 ratio (based on the cell count) with ^15^N-labelled *B. subtilis* PY79 vegetative cells. After mixing, samples were stored in 20% methanol at −20□ after freezing in liquid nitrogen. To quench further protein synthesis, 100μg/ml chloramphenicol was added to the mixture prior storage. This approach was used for the experimental analysis of spore revival.

#### 2.5.2. Approach II: ^13^C_6_, ^15^N_2_-Lysine incorporation to assess amino acid transporter activity

The *B. subtilis* PB2 ^14^N-spores were germinated in MOPS minimal medium with L-Lysine-^13^C_6_, ^15^N_2_ hydrochloride (Sigma Aldrich, henceforth referred as ^SILAC^Lysine) along with a mixture of AGFK and L- alanine as described previously. Samples were taken at t = 15, 45, 210 and 250 minutes.

#### 2.5.3 Approach III: Analysis amino acid recycling by spores

*B. subtilis* PY79 spores were labelled with heavy isotope of nitrogen (^15^N) and allowed to germinate in MOPS minimal medium supplemented with ^14^NH_4_Cl and a mixture of AGFK and L- alanine as described earlier. Samples for the dormant spores at t = - 30 minutes, for the spores after heat activation t= 0 and during germination at t =15, 90, and 150 minutes were harvested.

#### 2.5.4. Peptide isolation and fractionation

In Approach I, samples from were subjected to one pot isolation method and peptides were fractionated using ZIC-HILIC^5,13^. For approach II and III fractionation was omitted. Instead, the tryptic digest was freeze-dried before use and the freeze-dried samples were re-dissolved in 0.1% TFA and desalted using Omix μC18 pipette tips (80-μg capacity, Varian, Palo Alto, CA, USA) according to the manufacturer’s instructions.

#### 2.5.5 FT-ICR MS/MS

ZIC-HILIC fractions were re-dissolved in 0.1% TFA and peptide concentrations were determined by measuring the absorbance at wavelength of 215 nm with a NanoDrop. LC-MS/MS data were acquired with an Apex Ultra Fourier transform ion cyclotron resonance mass spectrometer (Bruker Daltonics, Bremen, Germany) equipped with a 7 T magnet and a Nano electrospray Apollo II Dual Source coupled to an Ultimate 3000 (Dionex, Sunnyvale, CA, USA) HPLC system. Samples containing up to 300 ng of the tryptic peptide mixtures were injected as a 10μl 0.1% TFA, 3% ACN aqueous solution together with 25 fmol of [Glu1]-Fibrinopeptide B human peptide and loaded onto a PepMap100 C18 (5 μm particle size, 100 Å pore size, 300 μm inner diameter × 5 mm length) precolumn. Following injection, the peptides were eluted through an Acclaim PepMap 100 C18 (3 μm particle size, 100 Å pore size, 75 μm inner diameter × 250 mm length) analytical column (Thermo Scientific, Etten-Leur, the Netherlands) to the Nano electrospray source. Gradient profiles of up to 120 min were used from 0.1% formic acid-3% acetonitrile to 0.1% formic acid-50% acetonitrile (flow rate 300 nl/min). Data-dependent Q-selected peptide ions were fragmented in the hexapole collision cell at an argon pressure of 6 × 10-6 mbar (measured at the ion gauge) and the fragment ions were detected in the ICR cell at a resolution of up to 60,000. In the MS/MS duty cycle, three different precursor peptide ions were selected from each survey MS. The MS/MS duty cycle time for one survey MS and three MS/MS acquisitions was approximately 2s. Instrument mass calibration was better than 1 ppm over m/z range of 250– 1500.

#### 2.5.6 Data analysis and bioinformatics

Raw data was analysed in a similar way as described in previously^5^. The consecutive time points were analysed by paired *t*-test to obtain *p*-value for each protein. Proteins with *p*-value < 0.05 were considered to be differentially expressed. The ^14^N/^15^N ratios obtained by Approach I were Z-transformed prior to K-mean clustering. DAVID Bioinformatics Resources tool (version 6.8) was used^19,20^ to retrieve the Uniprot key word enrichment and GO Term classifications. Identified proteins were categorized according to SubtiWiki (http://subtiwiki.uni-goettingen.de/)^21^.

## 3 Results

### 3.1 Core set of transcripts and proteins in a dormant spore

Extensive research in recent years has confirmed that the dormant spores retain transcripts throughout their dormancy^1,22-25^. We have identified 36 transcripts present abundantly in the dormant spores. This number is in close agreement with the previous findings^1,24^. These transcripts mainly belong to the genes encoding small acid soluble proteins (*sspE, sspO, sspI, sspK, sspF*) and proteins with uncharacterized functions.

Using Approach I, 1086 proteins are quantified, in at least two independent biological replicates, from the dormant spores. The Uniprot terms enriched from this dormant spore proteome are depicted in **Figure 2.** According to the DAVID functional enrichment, the proteins belonging to the functional classes of ribosome biogenesis, carbon metabolism, RNA processing and protein synthesis are highly enriched in the dormant spores.

**Figure 2.**
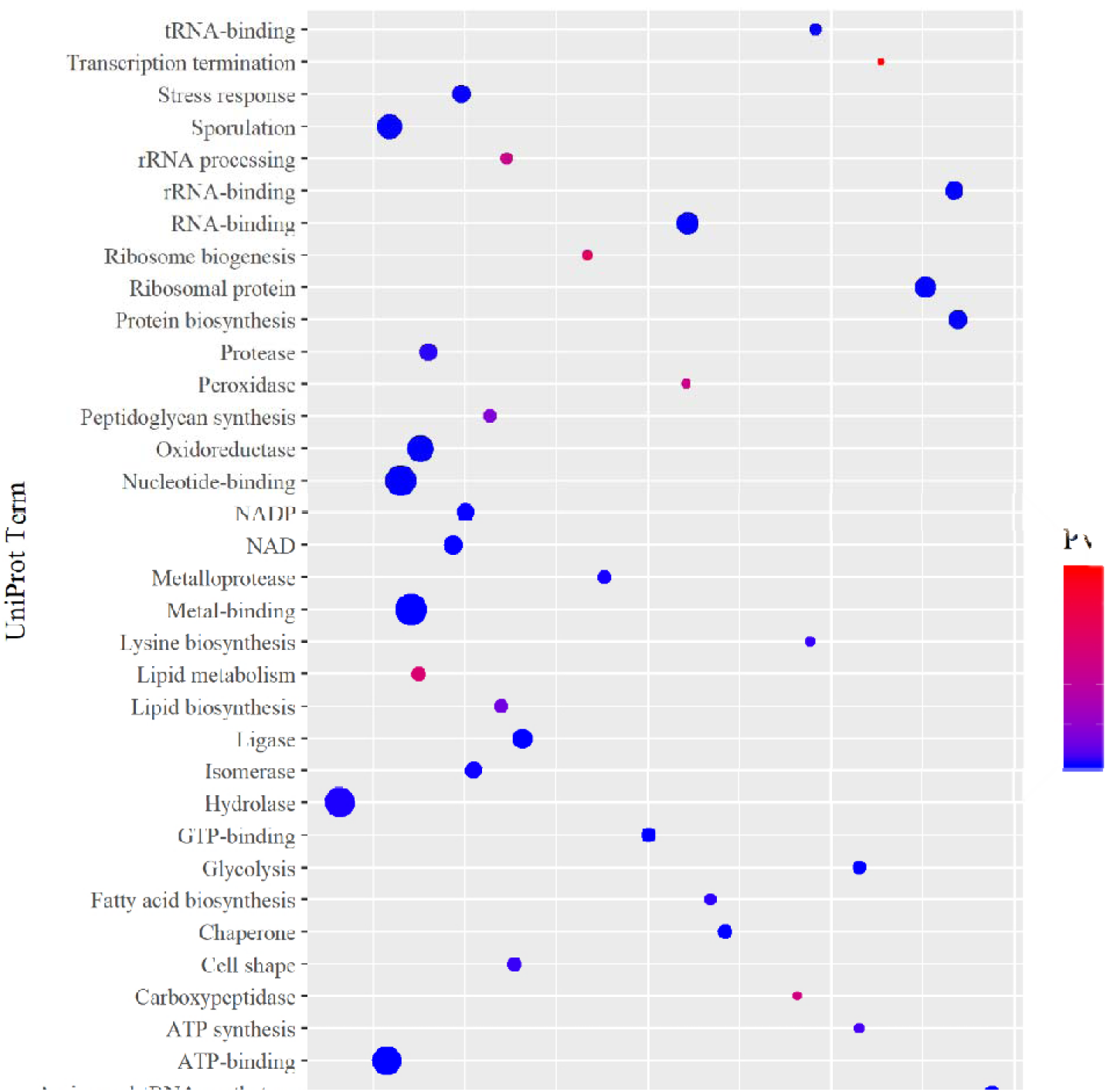
Uniprot categories enriched from the quantified proteins in the dormant spores by DAVID analysis. The fold enrichment is the fraction of quantified proteins belonging to a particular UniProt category as compared to the total number of proteins assigned to that category in the genome.

### 3.2 Comparative analysis of transcriptome and proteome

Integrated time-resolved analysis of mRNA gene-expression profiles and their resulting proteins may yield intriguing insights about the cellular level at which the molecular physiology of any organisms under study is mostly controlled. Our analyses show that during heat activation, there is no significant expression of transcripts but throughout the subsequent spore revival (outgrowth) phases, 3152 differentially expressed transcripts have been observed based on the analysis approach described in the Methods and Methods section using an adjusted *p*-value < 0.01. These transcripts are divided into 40 clusters by K-means clustering with transcripts of different functional categories showing similar expression profiles per cluster (**Supplementary Figure 1, Supplementary Table 1**). In case of proteome changeover analysis, 773 proteins have been quantified across the germination time line in at least two replicates (**Supplementary Table 2**). Of these 451 proteins are present in all the replicates and at all the time points. These proteins are further considered for data analysis. It is evident that 323 of these proteins are differentially expressed (DEPs) while the levels of 128 proteins remain relatively stable. K-means clustering of DEPs led to 10 clusters (**Figure 3)**. Cluster 1 consists of ABC transporters, aminoacyl tRNA synthetases along with chaperones. The relative levels of these proteins do not change significantly till 60 min after germination. On the contrary, the proteins involved in amino acid biosynthesis (cluster 4) show a steady increase immediately after germination. In cluster 5, significant change in relative protein expression is observed at later stages of outgrowth (t = 210). This group comprises of, ribosomal proteins, proteins involved in the TCA cycle as well as chemotaxis proteins. Interestingly in cluster 6, protein expression profiles show two stages. Here the first increase in relative levels of proteins is observed between 30 to 60 min of germination and the second increase is at the time of burst (t = 150). Purine biosynthetic proteins, protein involved in DNA replication, cell division and septation proteins show such clustered profile. In clusters 1, 3 and 10, proteins decrease in their relative levels initially during germination. These clusters involve proteins belonging to few aminoacyl tRNA synthetase and glycolysis. Proteins from clusters 2 show a scattered trend and these proteins are mainly spore specific uncharacterized proteins.

**Figure 3.**
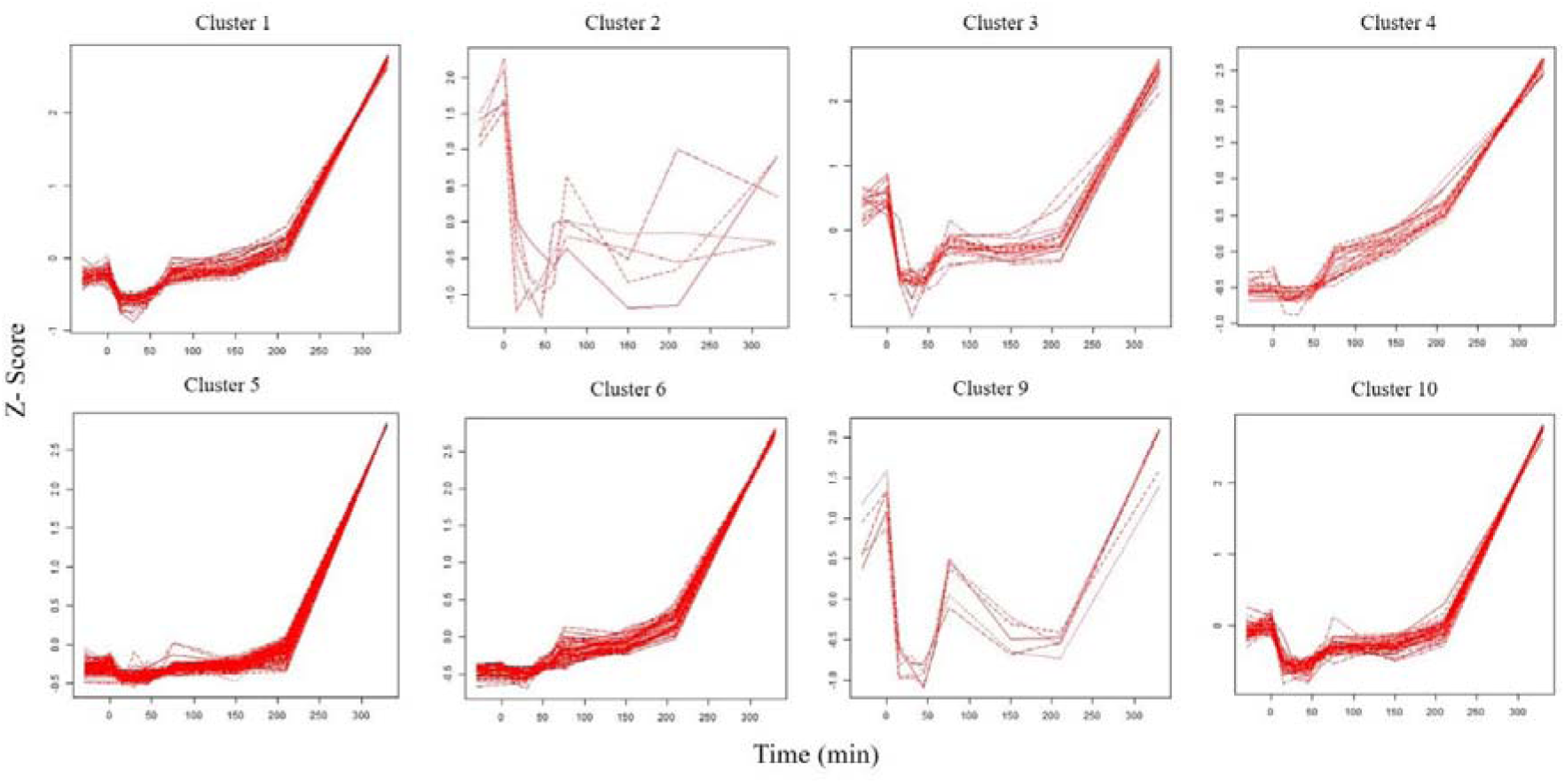
Temporal clustering during spore germination and outgrowth. The differentially expressed proteins are organized into 10 clusters by K-means clustering. Clusters 7 and 8 are not included as they consist of less than four proteins.

### 3.3 Differentially expressed genes (DEGs) and proteins (DEPs)

Figure 4 shows a heat map of differentially expressed genes belonging to five functional classes and the relative levels of the corresponding proteins at different time points. At the start of germination, (0 to 15 min) among others, 14 transcripts related to purine and pyrimidine biosynthesis, 9 transcripts associated with amino acid biosynthesis and 31 transcripts belonging to the translational machinery are upregulated. Additionally, seven genes belonging to the central metabolic pathways are also slightly upregulated in this period. The levels of the analogous proteins encoded by all these transcripts increase post 30 minutes of germination (**Supplementary Table 3 (A)**).

**Figure 4.**
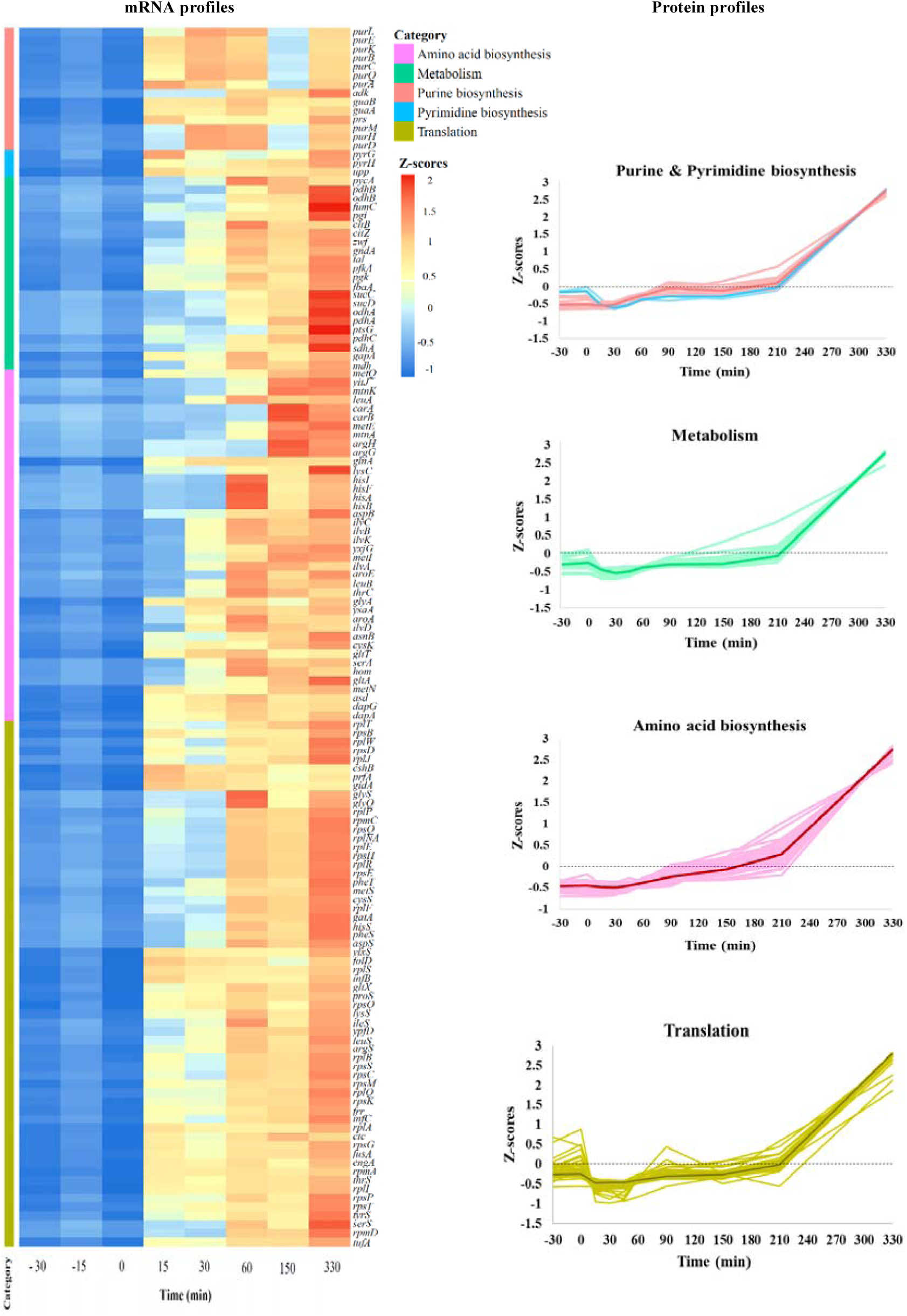
Functional categories and expression patterns of the differentially expressed genes and proteins during germination and outgrowth of *B. subtilis* spores. The Z-score transformed profiles of the genes (rows in the heatmap) and corresponding proteins are shown. The columns represent the different time points from dormant to outgrowing spore. The light colours in the protein profiles correspond to the individual proteins whereas the dark colours indicate the median Z-score profiles. Functional categories are obtained from SubtiWiki^21^.

Transcription of genes contributing to the DNA replication, processing and repair, peptidoglycan and fatty acid biosynthesis, cell division and cell shape also initiates during phase transitioning **(Figure 5, Supplementary Table 1)**. As seen, rather a minority of proteins from these categories are seen as differentially expressed. In contrast, for the functional groups, transporters and membrane proteins, large proportion of proteins are subjected to temporal variation. These include ATP synthesis and hydrolysis proteins (e.g AtpA, AtpD), protein transporters (e.g SecDF, OppA), proteins involved in glycolysis and TCA (e.g GapA, Eno, PdhABCD), iron transporters (e.g YfiY, YhfQ), amino acid transporters (e.g GltT, MetN) as well as some permeases and ion transporters (**Figure 5, Supplementary Table 3 (A)**).

**Figure 5.**
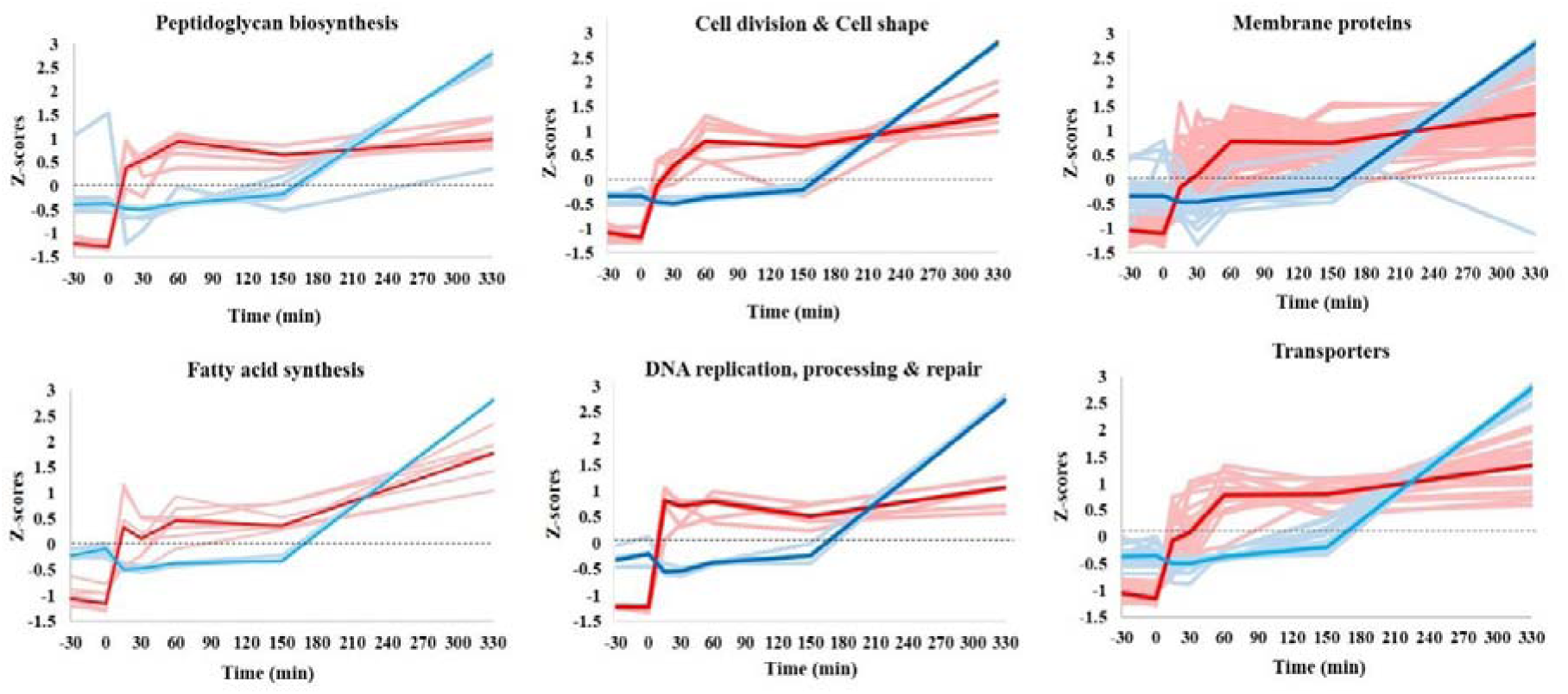
Representation of the trends in mRNA expression and relative changes in protein expression in germinating spores of *B. subtilis.* The trends for the differentially expressed mRNA and proteins belonging to the functional categories peptidoglycan biosynthesis (n = 8), fatty acid synthesis (n = 8), cell division and cell shape (n = 7), DNA replication, processing and repair (n = 5), membrane proteins and transporters (n = 69, 21 respectively) are shown. The light colour represent the individual genes (red) and proteins (blue) whereas the dark colour indicates the median Z-scores for the set of genes and proteins across all the time points. The trends for proteins represent the changes in the protein levels relative to those in the vegetative cells. Functional categories are obtained from SubtiWiki^21^. Refer to **Supplementary Table 4.3** for more details.

#### 4.3.4 Non-Differentially expressed proteins (NDEPs)

There are 128 proteins, quantified in all three replicates at all the time points (**Supplementary Table 3(B)**), which do not show any significant variation in their expression. Among these non-differentially expressed proteins, 12% belong to the category of proteins related to sporulation and spore structure. These include inner coat proteins such as SpoIVA, SafA, CotI, CotS, CotSA, outer coat protein CotB and glycolytic proteins Eno, Pgm that are conserved in spores. Proteins involved in ribosome biogenesis (CshA & ObgE) along with some ribosomal proteins like RpsJ, RplK and RpmE2 also seem to be unaffected during germination indicating the adequacy of their levels in the dormant spores. Amino acid transporter proteins TcyA, OpuCC, ArtP also seem to remain stable in their expression patterns. Ion transporters AtpC and the important germination related calcium transporter AtcL (YloB) also maintain stable expression levels.

The PTS fructose transporter FruA, present abundantly in the dormant spores, is maintained relatively stable until the outgrowth phase. Such behavior is also common with methyl accepting chemotaxis proteins McpC and TlpB. Thirteen proteins with unknown function show relatively stable expression throughout the spore revival (**Supplementary Table 3(B)**). The spore-associated protein YodI falls in this group.

#### 4.3.5 Signs of protein synthesis

In order to gain more insights in the protein synthesis during germination, two alternative experimental strategies have been explored in addition to Approach I. Firstly, protein synthesis during germination has been monitored by germinating the ^15^N-metabolically labelled PY79 spores in ^14^N minimal medium (Approach III). Individual LC-MS spectra from the samples have been scanned for ^14^N incorporation of tryptic peptides in newly synthesized proteins (**Figure 6**). Secondly, protein synthesis has also been monitored by germinating ^14^N PB2 spores in ^14^N minimal medium including ^SILAC^Lysine (Approach II). Incorporation of ^SILAC^Lysine introduces a mass shift of 8 Da for each of the respective ^14^N peptide. The newly synthesized proteins have thus been quantified by calculating the [^SILAC^Lysine] / ^14^N protein ratios. These ratios are included in the **Supplementary Table 4.** The incorporation of ^SILAC^Lysine and recycling of ^15^N amino acids and (Approach II and III) are both visible in a number of proteins during and immediately after germination. In the first 15 minutes past germination, 20 proteins show this incorporation whereas this number increases to 119 in the next 30 minutes **(Supplementary Table 4). Figure 6** shows two examples of such proteins. Panel **A1** shows the germination profile (obtained by Approach I) of the triply charged SVDPAANPYLALSVLLAAGLDGIKNK tryptic peptide from glutamine synthetase relative to the corresponding ^15^N peptide of the reference vegetative cells. The synthesis of this peptide starts around 15 minutes after addition of germinants. After a gradual increase in the peptide levels for 90 minutes, the peptide synthesis appears to slow down until 200 minutes. Later, it speeds up again to facilitate the spore’s outgrowth further and prepare it for progression to the first division. All quantified tryptic peptides from glutamine synthetase show the same time profile (Cluster 6).When ^15^N-labelled spores are germinated in ^14^N germination medium (Approach III), the LC-MS spectra of the same peptide show minimal amount of the protein in dormant spores (t= −30) and there is no change until after heat activation (t = 0). Also in this analysis, the onset of protein synthesis is seen around 15 minutes after germination initiation. Significant increase of the ^15^N peptide level at this point implies recycling of the ^15^N amino acids. In addition, incorporation of ^14^N amino acids after 15 minutes results in a peptide with a mix of ^14^N and ^15^N amino acids. This indicates that synthesis of ^14^N amino acid co-occurs with the recycling of ^15^N amino acids. During the ripening outgrowth phase (90-150 minutes) the ^15^N recycling and ^14^N amino acid synthesis both progress steadily and after 150 minutes, the recycled ^15^N amino acids appear to run out while more and more ^14^N amino acids are incorporated. For evaluation, the simulated isotope pattern of the pure ^14^N tryptic peptide is shown (in blue, **Figure 6. A2)** in the MS spectrum. Incorporation of ^SILAC^Lysine in the triply charged SVDPAANPYLALSVLLAAGLDGIKNK tryptic peptide is observed 15 minutes after germination (Approach II). After 45 minutes both ^14^N and ^SILAC^Lysine peptide levels have increased, implying that recycling of ^SILAC^Lysine and incorporation of ^SILAC^Lysine both occur. After 190 minutes only the ^SILAC^Lysine peptide level has increased, implying that the spore runs out of ^14^N recycled lysine. After 250 minutes the ^SILAC^Lysine peptide level has increased further (**Figure 6. A3**). For the 30S ribosomal protein S1 (YpfD), its relative levels in the dormant spores (t = −30 min) are about 20% compared to those in the vegetative cells. Slight increase of the QSGIIPISELSSLHVEK tryptic peptide of this protein after about 90 minutes, indicates the onset of protein synthesis about 90 minutes after germination. As for the glutamine synthetase, the protein synthesis speeds up after about 200 minutes (**Figure 6. B1**). All quantified tryptic peptides from YpfD show exactly this time profile (Cluster 1). For the ^15^N labelled spores germinated in ^14^N medium, the LC-MS spectra show no significant increase in the levels of 30S ribosomal protein S1 until 15, 90 and 150 minutes. After 90 minutes peptides appear with a mix of ^14^N and ^15^N amino acids, showing that synthesis has started using both recycled ^15^N and newly synthesized ^14^N amino acids (Approach III). After 150 minutes, synthesis has hardly progressed, in agreement with the protein level profile in (**Figure 6. B1**). After 45 minutes incorporation of ^SILAC^Lysine discreetly appears, while the ^14^N peptide level is unchanged. After 210 and 250 minutes, the ^SILAC^Lysine peptide level has modestly increased (**Figure 6. B3**), in agreement with the protein level profile in **Figure 6. B1** and the ^14^N amino acid incorporation in **Figure 6. B2**.

**Figure 6.**
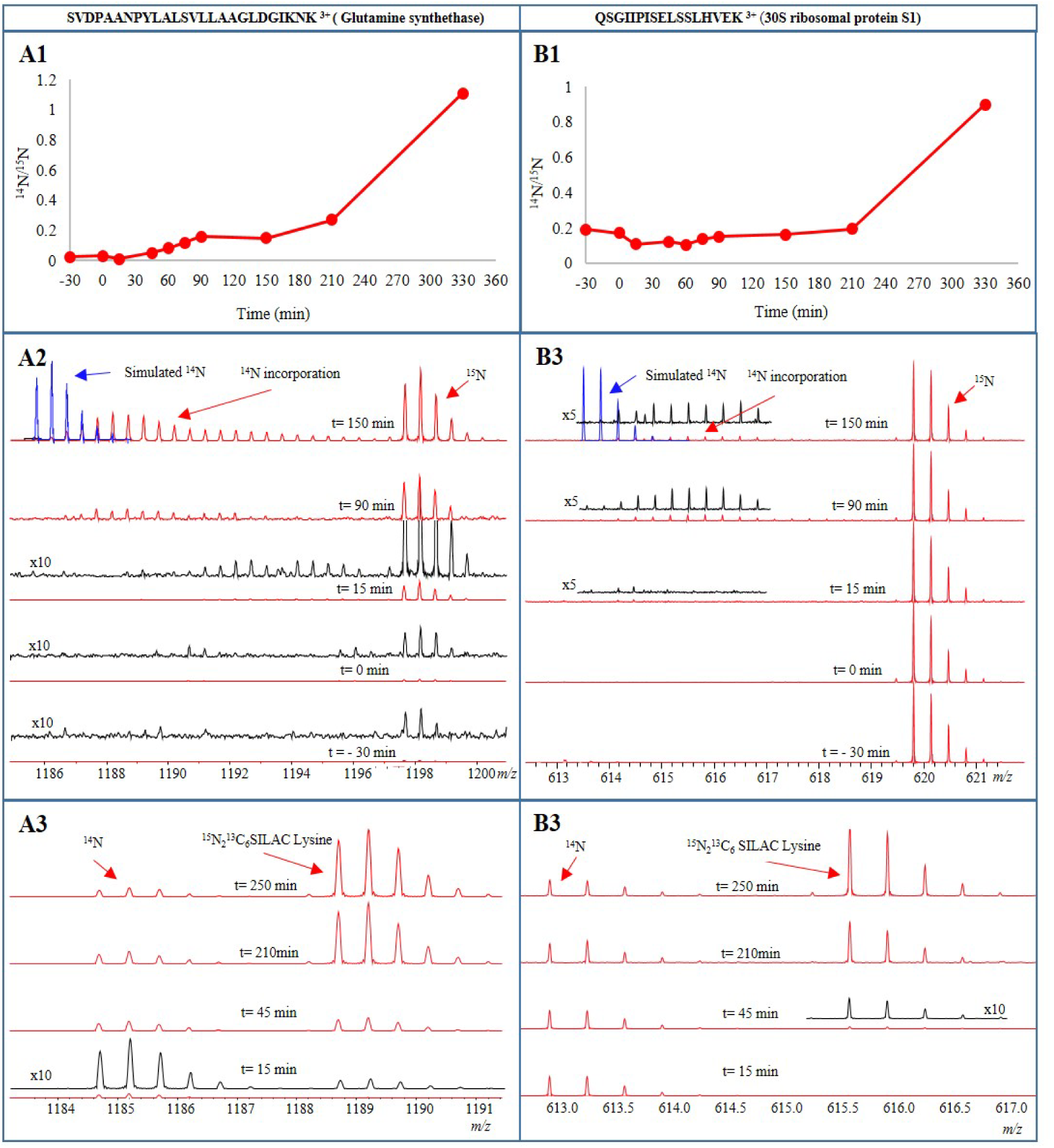
The A and B panels correspond to the time profiles of the triply charged tryptic peptide from Glutamine synthetase and the triply charged tryptic peptide from 30S ribosomal protein S1, respectively. **A1** and **B1**: The ^14^N peptide profiles relative to the corresponding ^15^N peptide from the reference ^15^N labelled vegetative cells. **A2** and **B2**: Incorporation of ^14^N amino acids in the tryptic peptides during synthesis of the corresponding proteins in ^15^N labelled spores during germination in ^14^N germination medium. **A3** and **B3**: Incorporation of ^SILAC^lysine in the tryptic peptides during synthesis of the corresponding proteins in ^14^N spores during germination in ^14^N germination medium with ^SILAC^lysine. For better visualization, MS spectrum is zoomed five (5X) and ten (10X) times.

## 4. Discussion

For bacilli and clostridia, spore germination is a highly efficient and systematic means to revert from the metabolically dormant state, the spore, to which these bacteria adhere upon exposure to severe environmental stress conditions. However, a significant heterogeneity is observed in germination of spores within a single isogenic population. In the last decade, the rationale behind this heterogeneity has been studied extensively and some preliminary putative molecular mechanisms that may be at the basis of this observation have been identified^26-28^. Recently, the aspect of mRNA and protein synthesis in a germinating spore has been re-discussed after a gap of many years. To this extent, some studies have shown the mRNA as well as protein expression profiles of germinating *B. subtilis* spores and yet the dilemma about the necessity of protein synthesis for spore germination remained unsettled. In the present study, we aimed to analyze the mRNA and protein expression patterns genome wide in an integrated manner. Thus, for the first time, in *Bacillus subtilis* an integrated view of its transcriptome and quantitative proteome upon spore germination and outgrowth emerged. In order to perform such comparative analysis we have germinated the spores in a minimal medium supplied with germinants, which leads to a moderate speed of progression of the germination and outgrowth process^29^ enabling adequate sampling for the foreseen time resolved mRNA and protein studies.

It is well known that during the initial phases of spore germination, the water from the environment enters the dehydrated spore core thereby reactivating the latent processes within the spore. Characteristically, the mRNA and protein synthesis are the primary processes to be activated as was already demonstrated long time ago by Ginsberg and Keynan^30^. In agreement with this observation, our data confirms that the transcription machinery is highly active during and beyond the phase transition period with the expression of ∼ 2400 mRNA transcripts being upregulated. This is necessary as for many transcripts <1 molecule is present in a dormant spore^24^. Therefore, along with transporters^1^, the genes related to the processes of transcription regulation, translation, DNA replication and repair, rRNA processing, ribosome as well as inosine and uridine monophosphate (IMP-UMP) biosynthesis, cell shape and cell division are evidently upregulated in this first phase of transcription initiation. At the same time, some spore transcripts as well as the transcripts of spore associated hypothetical protein genes are seen to be downregulated (broken down) which may serve as the source of nucleotides (for the *de novo* nucleotide synthesis) as suggested previously^24^. In the next phase of transcription, genes belonging to the amino acid biosynthesis, protein/peptide transport are upregulated, which may later facilitate the transport of newly synthesized proteins and peptides. As the ripening phase stabilizes, genes related to glycolysis, the TCA cycle, branched chain and aromatic amino acid biosynthesis are activated. Note that many of the proteins belonging to these basic metabolic pathways are actually resident spore proteins^5^. Hence, they are part of the ‘survival kit’ for bacterial life, are not in need of early synthesis during germination or outgrowth and in fact can thus mediate the metabolic requirements at the onset as well as during spore revival. Further, in later stages of outgrowth, the outgrowing spore appears to prepare for the ‘*burst****’*** as it synthesizes enzymes involved in the biosynthesis of main cell envelope macromolecules^31^. In order to equip itself with sufficient nitrogen and sulphur stores, the expression of genes responsible for the urea cycle and hydrogen sulfide production is triggered. Ultimately, as the outgrowth is about to complete, the spore has its DNA duplicated, amino acids synthesized, transcription and translation active, metabolism restored, transporters activated and proteins available to carry out the cell elongation and division. Thus the highlight of the final phase of transcription are the genes responsible for the cell division and cell cycle proteins, chemotaxis and ion homeostasis proteins, stress response proteins as well as the flagellar assembly proteins. Remarkably, the purine and pyrimidine biosynthesis processes appear to progress in a two-step manner. The initial rise in the transcripts takes place in the first 15 minutes while the second increase in the transcript levels is around the burst time. Such behaviour corroborates old observations made in *B. cereus* spores^32^. Still, the reason why this two-step increase is seen remains to be elucidated.

Similar to the mRNA synthesis, the protein synthesis in spores also progresses as groups of (functionally) related proteins appearing simultaneously alluding to the occurrence of a germination protein synthesis program. The phase transition period is characterised by loss of about 30% of the spore dry weight^33^, during which many proteins and peptides are lost along with the CaDPA. Our data shows that 173 proteins from the dormant spores, some being from the shared category^5^, are downregulated during the phase transition period. This loss may partially be ascribed to the active protein degradation, undertaken by the spore proteases such as Gpr, as germination proceeds. Active degradation of SASP proteins by Gpr is well studied^34^. Our data clearly exemplifies this sequential break down for SspE (**Supplementary figure 2**). Such initial protein degradation leads to an additional pool of free amino acids which are recycled for the renewed protein synthesis^10^. Such early signs of protein synthesis are clearly visible in our data (**Figure 6**). It is remarkable that abundant ^SILAC^Lysine incorporation or ^15^N amino acid recycling coincides with the fast degradation of small acid soluble proteins (SASPs). Amongst the early synthesized proteins, those involved in purine and pyrimidine biosynthesis are present. Although a total of 25 proteins belonging to these categories have been quantified at all the time points in our data, only 17 are seen to be differentially expressed throughout the germination and outgrowth time series (**Supplementary Table 3**). Surprisingly the pyrimidine biosynthesis proteins are significantly upregulated only at the end of the germination time line. These observations are in synchrony with the respective gene transcription events. At the same time the proteins required for isoleucine and valine biosynthesis, glutamine as well as homocysteine metabolism are triggered in order to initiate amino acid biosynthesis in the next phase of translation. After these initial preparations (> 30 min), the spore carries out translation of a number of proteins that are central to glycolysis, gluconeogenesis, amino acid biosynthesis, AMP synthesis and acetyl-CoA synthesis. The chromosomal replication initiator protein DnaA is seen to be synthesized in this period and its levels remain constant through outgrowth. Notably, the proteins belonging to the histidine biosynthesis pathway are also induced which correlates with the synthesis of histidine tRNA ligase 15 to 30 minutes into outgrowth. In the period of 60 - 90 minutes post-germination, the spore shows bulk synthesis of ribosomes, as well as GTP-GMP biosynthesis proteins. Interestingly, the next phase of translation, when the spore is near to its burst time, is marked by the synthesis of sulfur-containing amino acids methionine and cysteine. The reactions involved are carried out by MetI and MetC proteins that are synthesized in this period. In addition, peptidoglycan biosynthesis proteins and those related with flagellar assembly are also synthesized prior to burst. Thus the spore ripening stage seems to be of paramount importance in the translational schedule of a germinating spore where proteins with varied functional aspects are synthesized, likely the actual outgrowth gene-expression program chosen is to a significant extent dependent on the is to a significant extent dependent on the is to a significant extent dependent on the is to a significant extent dependent on the environmental conditions encountered. Ultimately, when the spore breaks its dormancy and grows to become a vegetative cell, it is equipped with the transcription and translation machinery, as well as its basic energy metabolism activated. Finally, the proteins for the TCA cycle, ATP synthesis, mRNA processing, fatty acid synthesis and cell division are synthesized in a second stage. The inferred mechanistic strategy of the germinating and outgrowing spore in its quest for a return to (microbial) life is summarized in **Figure 7**. Throughout the germination and outgrowth alanine dehydrogenase (Ald), cell wall associated protein YoeB and transition state regulatory protein AbrB show continued increase in their relative levels compared to the vegetative cells. Conversion of alanine (provided as a germinant and also available as a free amino acid) to pyruvate by Ald may be a key reaction to form acetate^1,35^, ethanol and acetaldehyde. Especially acetate formation is an energy producing step^36^ that is beneficial for germination and outgrowth. It has been observed that pyruvate formed via Ald may be converted to fructose-6-phosphate via gluconeogenesis thus fuelling cellular energy generation processes^37^. Protein YoeB (IseA), an autolysis activation modulator^38,39^ may serve as a controller restricting cell division ahead of time. Similarly, the master regulator AbrB that suppresses transcription of many sporulation specific genes during vegetative growth increases significantly at the time where the spore is preparing for the burst. Thus it is speculated that, prior to the emergence of the first cell, the spore attains an optimum energy level, restricts untimely cell division and premature sporulation through intricate networks of signalling molecules that steer stress surveillance and response mechanisms.

**Figure 7.**
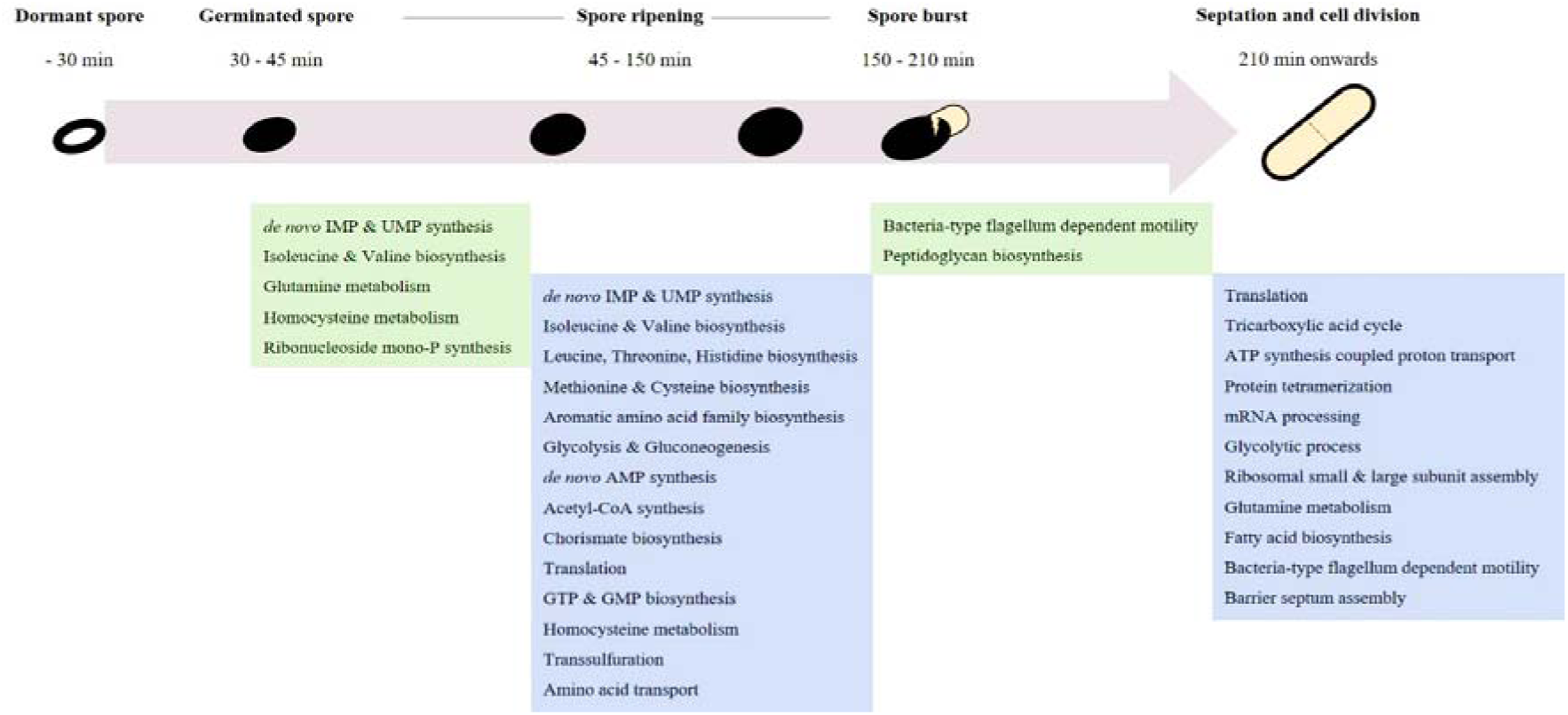
Summary of GO Terms enriched from the differentially expressed proteins at respective time points during *B. subtilis* spore revival in minimal medium.

In order to understand the spore’s putative germination plan, time resolved studies are highly effective. Especially to study the protein turnover during germination an accurate and sensitive method is a pre. Using ^15^N-metabolic labelling, we have successfully analysed the trend in overall protein dynamics at various time points during germination and outgrowth. However, these turnover profiles are always relative to the protein levels in the reference vegetative cells. Therefore, for spore specific and structural proteins, which are not present in the reference vegetative cells, the changes in their levels cannot be estimated accurately. We circumvented this with an approach where we started with ^15^N ^13^C labelled lysine in a SILAC approach complemented by ^15^NH_4_Cl labelled spores.

Some dormant spore proteins, such as those related to amino acid or purine/pyrimidine biosynthesis, are shared between the spores and cells. These proteins are present at levels <5% of those in the vegetative cells and minute changes in their levels are prominent and accurately quantified throughout germination. Yet, for some ribosomal protein (RpsB, RpsE), chaperons (GroL, DnaK) and glycolytic proteins (Eno, Gpm) and proteins that are present at levels of 10-30% (as compared to their levels in the cells) slight modifications cannot be estimated. The SILAC data therefore helps in such cases and clearly shows that synthesis of these proteins, indicated by lysine incorporation, starts earlier after phase transitioning is completed (15 - 45 minutes). For instance, trigger factor (Tig), crucial in the analyses of Sinai *et al.*^3^ and elongation factor (Tsf) show 7-8% incorporation of the ^SILAC^Lysine in at least two quantified peptides (see **Supplementary Table 4**). In the ^15^N-metabolic labelling time series however, an escalation in their levels is only seen at later time points. Thus, the more sensitive SILAC approach indicates the onset of protein synthesis while the ^15^N-metabolic labelling approach indicates that irrespective of their synthesis the levels of those proteins remain relatively stable. Clearly, our data are in that sense way more comprehensive than the data by Sinai *et al.*^3^ who have not addressed the spore protein composition nor spore protein dynamics during germination and outgrowth. Based on our results we speculate that a limited protein synthesis is applied upon germination (phase transition) by the spore until it reaches the burst time. In addition, it is noteworthy that in the initial heat activation a few proteins are downregulated (**Supplementary Table 3**). This observation is parallel to the general notion that protein denaturation and/or degradation may take place during the heat activation step^40^.

Summarizing, in our experimental setup no protein synthesis occurs in a dormant spore and as soon as germination is triggered both transcription and translation processes are activated. There is a loss of proteins during the phase transitioning period but it is not clear whether this is a result of active protein degradation or an actual physical loss of these proteins (i.e. exudate). As opposed to a large number of DEGs, the number of DEPs is limited. This discrepancy demands further attention. Moreover, the molecular details about the role of heat activation and the dynamic interplay of mRNA and proteins during phase transitioning remain the topics for the future research.

