## Supplementary Figure 1 and 2 for "Integrative analysis of proteome and transcriptome dynamics during *Bacillus subtilis* spore revival"

**Supplementary Figures**

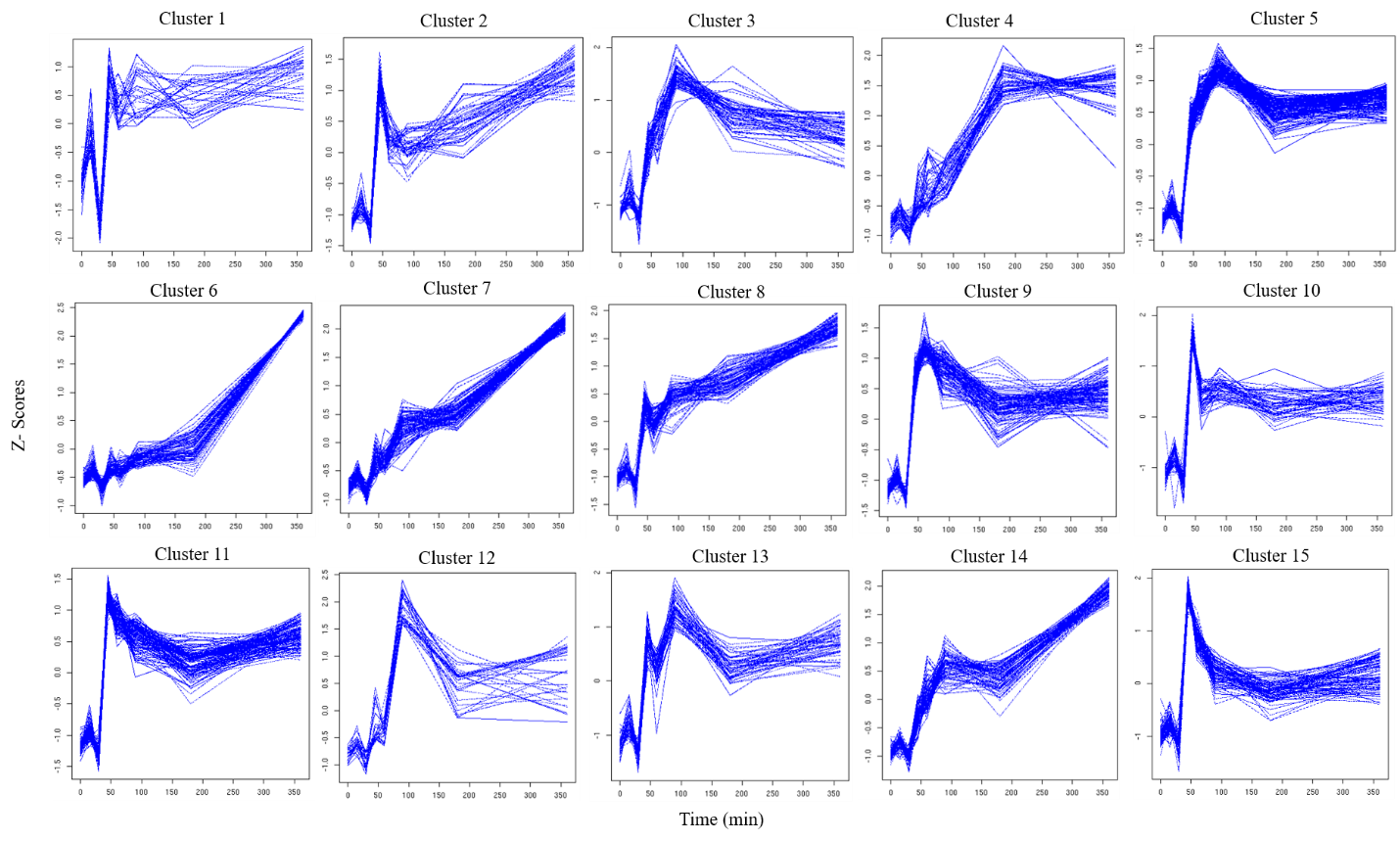
**Supplementary Figure 1: The K-mean clusters of mRNA transcription profiles of reviving *B. subtilis* spores.** K-mean clustering of the differentially expressed genes (DEGs) led to 40 clusters.

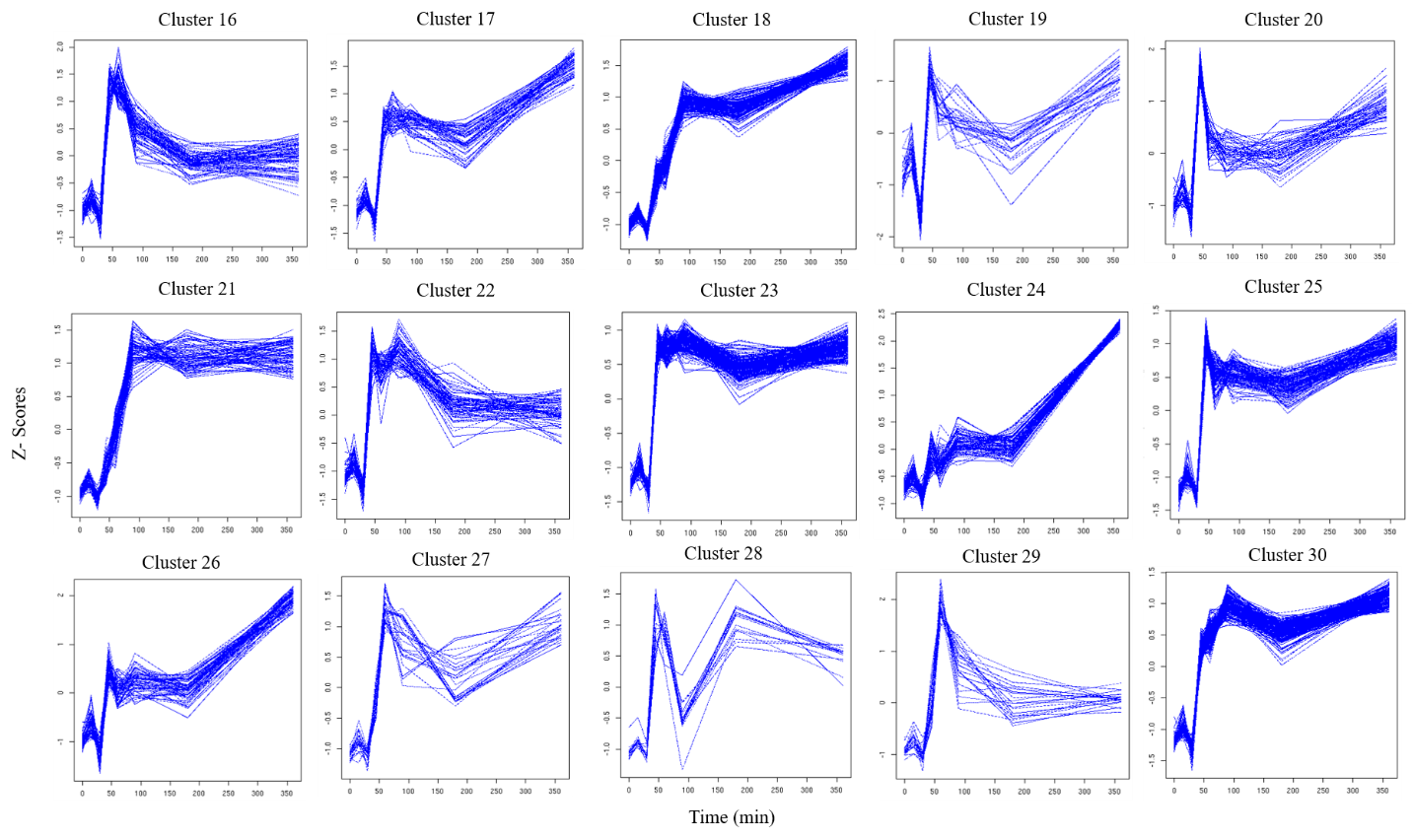

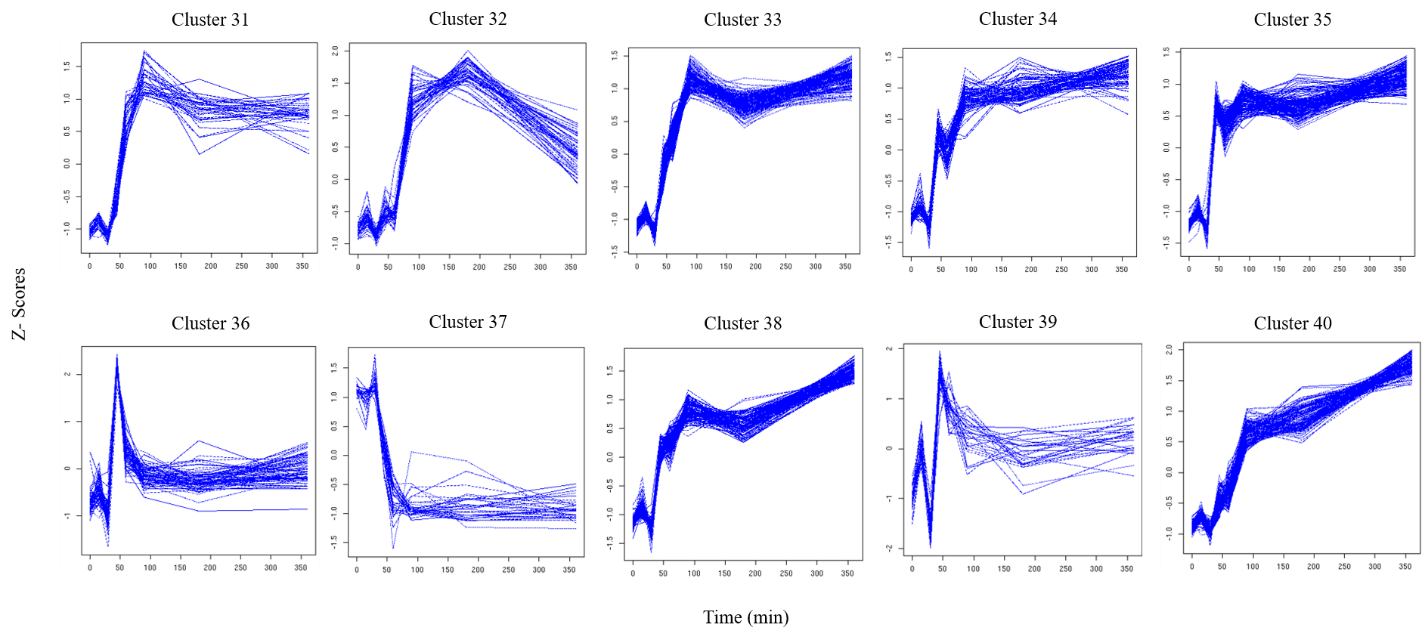

**Supplementary Figure 2: Sequential degradation of small acid soluble proteins (SspE and SspB) by Gpr.** The above peptide sequences have been identified in our analysis.

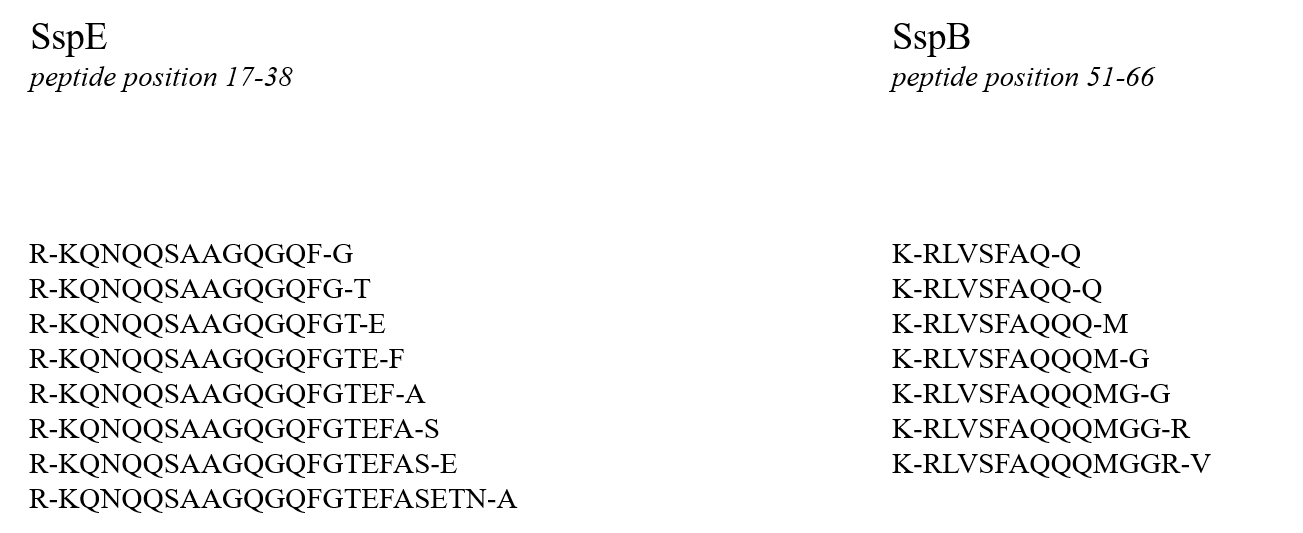
